# Refinement of a Cryo-EM Structure of hERG: Bridging Structure and Function

**DOI:** 10.1101/2020.09.11.293266

**Authors:** H.M. Khan, J. Guo, H.J. Duff, D. P. Tieleman, S. Y. Noskov

**Affiliations:** University of Calgary

## Abstract

The human *ether-a-go-go*-related gene (hERG) encodes the voltage gated potassium channel (KCNH2 or Kv11.1, commonly known as hERG). This channel plays a pivotal role in the stability of phase 3 repolarization of the cardiac action potential. Although a high-resolution cryo-EM structure is available for its depolarized (open) state, the structure surprisingly did not feature many functionally important interactions established by previous biochemical and electrophysiology experiments. Using Molecular Dynamics Flexible Fitting (MDFF), we refined the structure and recovered the missing functionally relevant salt bridges in hERG in its depolarized state. We also performed electrophysiology experiments to confirm the functional relevance of a novel salt bridge predicted by our refinement protocol. Our work shows how refinement of a high-resolution cryo-EM structure helps to bridge the existing gap between the structure and function in the voltage-sensing domain (VSD) of hERG.

**Statement of Significance:** Cryo-EM has emerged as a major breakthrough technique in structural biology of membrane proteins. However, even high-resolution Cryo-EM structures contain poor side chain conformations and interatomic clashes. A high-resolution cryo-EM structure of hERG1 has been solved in the depolarized (open) state. The state captured by Cryo-EM surprisingly did not feature many functionally important interactions established by previous experiments. Molecular Dynamics Flexible Fitting (MDFF) used to enable refinement of the hERG1 channel structure in complex membrane environment re-establishing key functional interactions in the voltage sensing domain.

## 1. Introduction

Human *ether-a-go-go*-related gene (hERG) is responsible for encoding the voltage gated potassium channel (KCNH2 or Kv11.1, commonly known as hERG)(1). Inherited mutations or non-regulatory blockade of hERG channels leads to long QT syndrome, and hence can lead to a potentially lethal form of cardiac arrhythmia. Several drugs designed for other cellular targets have been found to non-specifically block the hERG channel in ventricular myocytes. Therefore, the establishment of the high-resolution structure of hERG in its active (drug-binding) conformation is essential for the rational design of drugs that do not bind to hERG and was the focus points for many research labs over the last decade(2). The specific salt bridges involved in the stabilization of open (depolarized) and closed states of the channel have been studied by several groups using an arsenal of experimental methods ranging from including electrophysiology studies combined with targeted mutagenesis (3-10) combined with tryptophan scanning (11) or use of fluorescent tags(12) and FRET-based techniques(13, 14). The lack of experimentally determined structures led to the development of a plethora of structural models based on various templates as well as kinetic models aimed at providing a structural basis to a multitude of functional studies(1, 3, 4, 8, 15). A pivotal event for understanding of structure function relations in hERG channel was the publication of a high-resolution cryo-EM structure by Wang and MacKinnon in 2017 (16). The hERG channel 3D structure was refined to 3.8 Å resolution in presumably the depolarized or open state, which is the most stable functional state in the absence of a transmembrane voltage. This structure contains the entire Trans-Membrane Domain (TMD) as well as a large portion of the intra-cellular domains (PAS and CNBD).

Although this structure is obtained at a high resolution, it also raised several questions about how the structure relates to the extensive body of experimental data on the characteristic salt bridges and other interactions identified in the depolarized state of the voltage-sensing domain (VSD). For example, several salt bridges involved in the direct stabilization of the VSD in its depolarized state (1, 3, 15, 17, 18) are missing in the structure deposited to the PDB (PDB id: 5VA2). This indicates that the solved cryo-EM structure may not represent a fully functional state and needs to be refined further to obtain a structure which directly corresponds to the functional open conformation of the channel.

Another striking feature of the resolved structure was its different, “domain-swapped” VSD packing topology against the Pore Domain (PD), in stark contrast to the “non-swapped” topology reported from structural studies of voltage-gated K^+^ channels from the Shaker family(19, 20). The structural differences in packing of the VSD against the PD are likely reflected in differences in the gating process in the hERG channel and other related channels with a non-swapped topology compared to Shaker channels (12, 21-23). Such structural arrangement has several functional implications. One of the critical features of VSD in a non-swapped topology channels such as hERG or HCN seems to be plasticity of the voltage sensor(23), highlighting that homology models of hERG based on Shaker channels will not be able to capture this feature. Using tryptophan mutagenesis scanning, Subbiah et al. proposed a loosely packed configuration of the VSD for the open state of the channel (11), with space between the S1-S4 helices and the PD. More recently, a break in the helical structure of S4 and its apparent plasticity were proposed to be important gating determinants for the HCN channel (24). The gating charge associated with hERG activation is also surprisingly small compared to the gating charge of the Kv1.2 channel, suggesting a different mechanism(5). MD simulations performed with the cryo-EM structure embedded in a lipid bilayer also highlighted a loosely packed VSD configuration(25), in line with the observation of Subbiah et al.(11) However, the VSD in the cryo-EM structure is tightly packed against the PD. Shi et al. also emphasized the importance of further refinement of this cryo-EM structure due to the uncertain positioning of D509, which acts as a proton sensor and stabilizes the open state(18). All in all, these works highlight the importance of further investigating the structure of hERG, starting from the existing hERG cryo-EM structure, to obtain a fully functional state.

Our main goal is to refine the existing structure to overcome these inconsistencies between the structural state captured in cryo-EM and the plethora of functional and modelling studies that have established key interactions in the VSD of hERG. We would like to address two fundamental questions:

1. Can the apparent plasticity of the VSD be reconciled with the state deposited into PDB captured by the original cryo-EM density map and yet capture the missing functionally relevant interactions in the deposited PDB structure?
2. Is the VSD in hERG channel truly loosely packed as hinted by current computational(21) and previously published experiment studies and especially data from tryptophan/mutagenesis scanning(11, 18)? If not, will it still allow significant VSD hydration and changes in helicity as proposed by other studies performed on the channels with non-swapped topology?

To address these questions, we refined the existing hERG structure using Molecular Dynamics Flexible Fitting (MDFF)(26). We performed structural refinement in different model environments (vacuum, implicit solvent, bilayer) to investigate the possible effect of more native-like conditions on the fitting procedure and the importance of explicit lipids in establishing state-specific interactions in the VSD. We find that our bilayer-fitted MDFF structure has the highest structural quality (based on comparison of the global model fit to the reference map and the reference structure), side chain quality and re-establishes the missing salt bridges in the original cryo-EM structure, supported by literature. In addition, this procedure predicted a novel salt bridge. We have tested this novel salt bridge prediction with mutagenesis and electrophysiological experiments to show that this interaction is functionally important for activation of the hERG channel. Our work offers a computational approach followed by electrophysiology experiments to bridge the gap between the structure and function of hERG, which can be used for other ion channels and transporters.

## 2. Methods

### Systems setup

The cryo-EM structure of hERG (PDB id: 5VA2) was our starting point. As the cryo-EM structure was missing some loops (for example, loops involving following residues: 434-451, 511-519, 578-582, 598-602), we used the cryo-EM structure with modelled loops of the open state from Perissinotti et al. as the target structure for the MDFF(21). This structure contained the missing loops built using ROSETTA loop modelling, minimized and briefly equilibrated to avoid steric clashes, and did not contain the Per-Arnt-Sim (PAS) domain (21). This structure was used to generate a map at 5Å resolution excluding atoms forming the nanodisk. We choose 5Å as the VSD portion was resolved at close to 5Å resolution(16). Then we used the Molecular Dynamics Flexible Fitting (MDFF) (26-28) technique for structure refinement. We extracted the starting structure for fitting from our multi-microsecond MD simulation where the protein was embedded in a bilayer mimicking native environment(25). We extracted the structure towards the end of the simulation where the VSD was in fully expanded configuration. Different environmental conditions: vacuum, generalized born implicit solvent (gbis) and bilayer were tested during fitting to check the importance of the presence of lipids during the fitting process. Vacuum and gbis systems were prepared using the MDFF plugin(29) in VMD(30). For the bilayer system, the protein was embedded in a POPC bilayer (380 lipids in total, 190 in each leaflet). For the bilayer system, 0.15 M KCl solution was used. CHARMM-GUI was used to prepare the bilayer system(31).

### Simulation protocol

Simulations were carried out using the CHARMM36m force field for proteins(32) and CHARMM36 force field for lipids(33). NAMD(34) (v. 2.13) was used to carry out all the simulations. The gbis model was used for implicit solvent simulations. For the bilayer system, POPC lipids and explicit TIP3P(35) waters were used. Cut-off and switching distances for nonbonded calculations were as follows: 9 and 10 Å (vacuum), 15 and 16 Å (implicit solvent), 10 and 12 Å (bilayer). Integration of equations of motion was done using a 1 fs timestep for vacuum and implicit solvent, 2 fs for the explicit bilayer system. Temperature coupling was applied to all the systems. Semi-isotropic pressure coupling was applied to the bilayer system to preserve the bilayer shape; using Langevin piston method(36) with an oscillation period of 50 fs and a damping timescale of 25 fs. Vacuum and implicit solvent simulations were performed at 300 K, whereas bilayer systems were performed at 310 K and 1 atm pressure. For the gbis system, an ion concentration of 0.1 M was used and a surface tension value of 0.005 kcal/mol/Å^2^ was used for the nonpolar solvation. The solvent dielectric was set to 80. In all cases, secondary structure restraints along with cis-restraints on peptide bonds and chirality restraints were applied to the protein structure during fitting. We used a gscale value of 0.3 for all the cases. For the bilayer system, the coupling of the structure to the map was done gradually and at 10 ns steps for Cα atoms, the backbone atoms and finally the heavy atoms.

### Analyses

Root Mean Square Deviation (RMSD) and Cross-Correlation Coefficient (CCC) were calculated with MDFF plugins in VMD. Together they provide information on the overall structural quality compared to the target structure and the target cryo-EM map, respectively. The CCC is calculated between the generated maps from the fitted structure over the course of fitting simulation with the target map(26). Final fitted structures were validated using MolProbity (37, 38) (PHENIX(39) implementation) and using all four chains. MolProbity provides information on the overall side-chain quality of the structure. Salt bridges were analyzed using the Salt Bridges plugin in VMD, using a oxygen-nitrogen distance cut-off value of 3.2 Å.

### Molecular Biology

Methods for site-directed mutagenesis have been previously reported(40, 41). Single- and double-mutant constructs of hERG were produced using conventional overlap PCR with primers synthesized by Sigma Genosys (Oakville, Ontario, Canada) and sequenced using Eurofins MWG Operon (Huntsville, AL). The hERG constructs were transfected into human embryonic kidney (HEK) 293 cells by the calcium phosphate method and cultured in Dulbecco’s modified Eagle’s medium supplemented with 10% fetal bovine serum and 1% penicillin and streptomycin (GIBCO, CA).

### Electrophysiology

The extracellular solution contained (in mM) NaCl 140, KCl 5.4, CaCl2 1, MgCl2 1, HEPES 5, glucose 5.5, and was kept at pH 7.4 with NaOH. Micropipettes were pulled from borosilicate glass capillary tubes on a programmable horizontal puller (Sutter Instruments, Novato, CA). The pipette solution contained the following: 10 mM KCl, 110 mM K-aspartate, 5 mM MgCl2, 5mM Na2ATP, 10 mM EGTA - ethylene glycol-bis(-aminoethyl ether)-N,N,N,N tetra-acetic acid, 5 mM HEPES, and 1mM CaCl2. The solution was adjusted to pH 7.2 with KOH. Standard patch-clamp methods were used to measure the whole cell currents of hERG mutants expressed in HEK 293 cells using the AXOPATCH 200B amplifier (Axon Instruments). The holding potential was -80 mV. The amplitudes of tail currents were measured when the voltage was returned to -100 mV after + 50 mV 1-second depolarization.

#### Voltage-dependence of activation

From a holding potential of -80 mV cells were depolarized for 1 second to a range of voltages from –100 to +50 mV followed by a step to -100 mV (or -50 mV see Results, 1s) to record the tail currents. The isochronal tail current-voltage plots were fit to a single Boltzmann function (1):

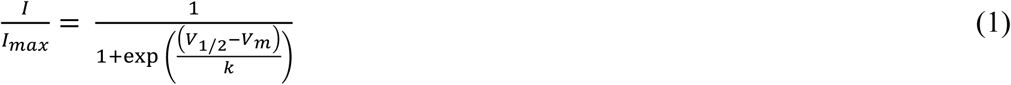

where *I/I*_*max*_ is the normalized current, V_1/2_ is voltage of the half-maximal activation, *k* is the slope factor and *V*_*m*_ is the membrane potential.

### Statistical Analysis of Electrophysiology data

The data are presented as the mean +/- S.D. One-way Anova test was used to analyse the data. P< 0.05 was designated as being significant.

## 3. Results

### 3.1 Environmental effects are important for the structure quality

The backbone RMSD with respect to the target structure is reported in Figure 1(A). The variation is minimal in the final segment of the trajectory for all three environments. This means that the final structures obtained using different environment during fitting converged to very similar configurations in terms of backbone RMSD. The obtained final structures retain very similar backbone configurations in the helical membrane parts of the protein (Figure 1(B)). There are small variations in the loop regions connecting the helices. The global CCC(26) values calculated over the course of fitting also converged to similar values (Figure 2). As can be seen, the final structures are very similar to the target structure in terms of their backbone configurations in the VSD and PD of the hERG. The resulting global CCCs from these structures are identical, indicating a high degree of global similarity to the target density map. These global comparisons highlight that we cannot distinguish which protocol performs better, either compared to the target structure or compared to the target map, as they converge to a very similar state.

**Figure 1.**
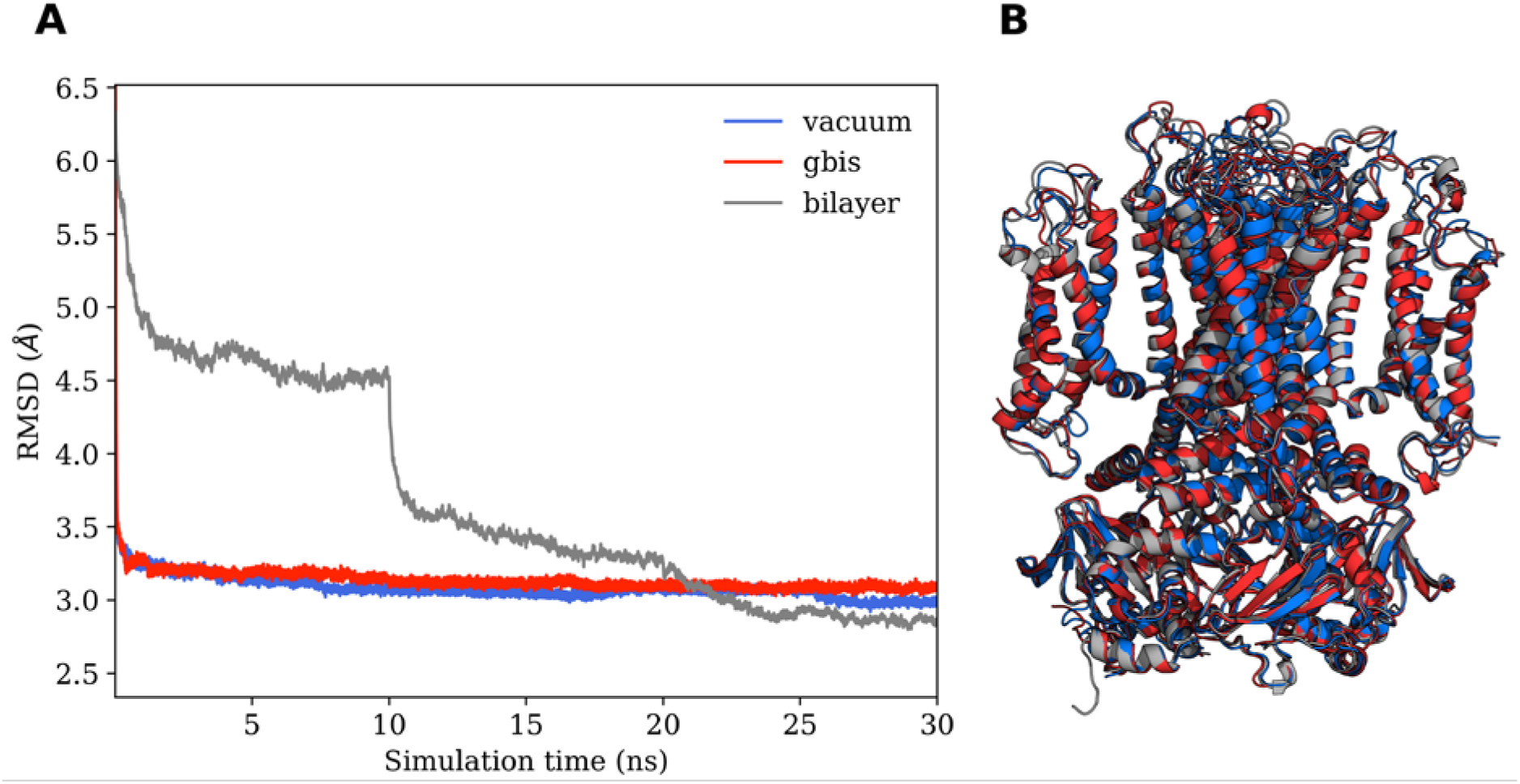
MDFF simulations performed in different environments. (A) Backbone RMSD of the protein during the course of the MDFF simulation, (B) final fitted structures obtained from the MDFF simulation: vacuum (blue), gbis (red), and bilayer (gray).

**Figure 2.**
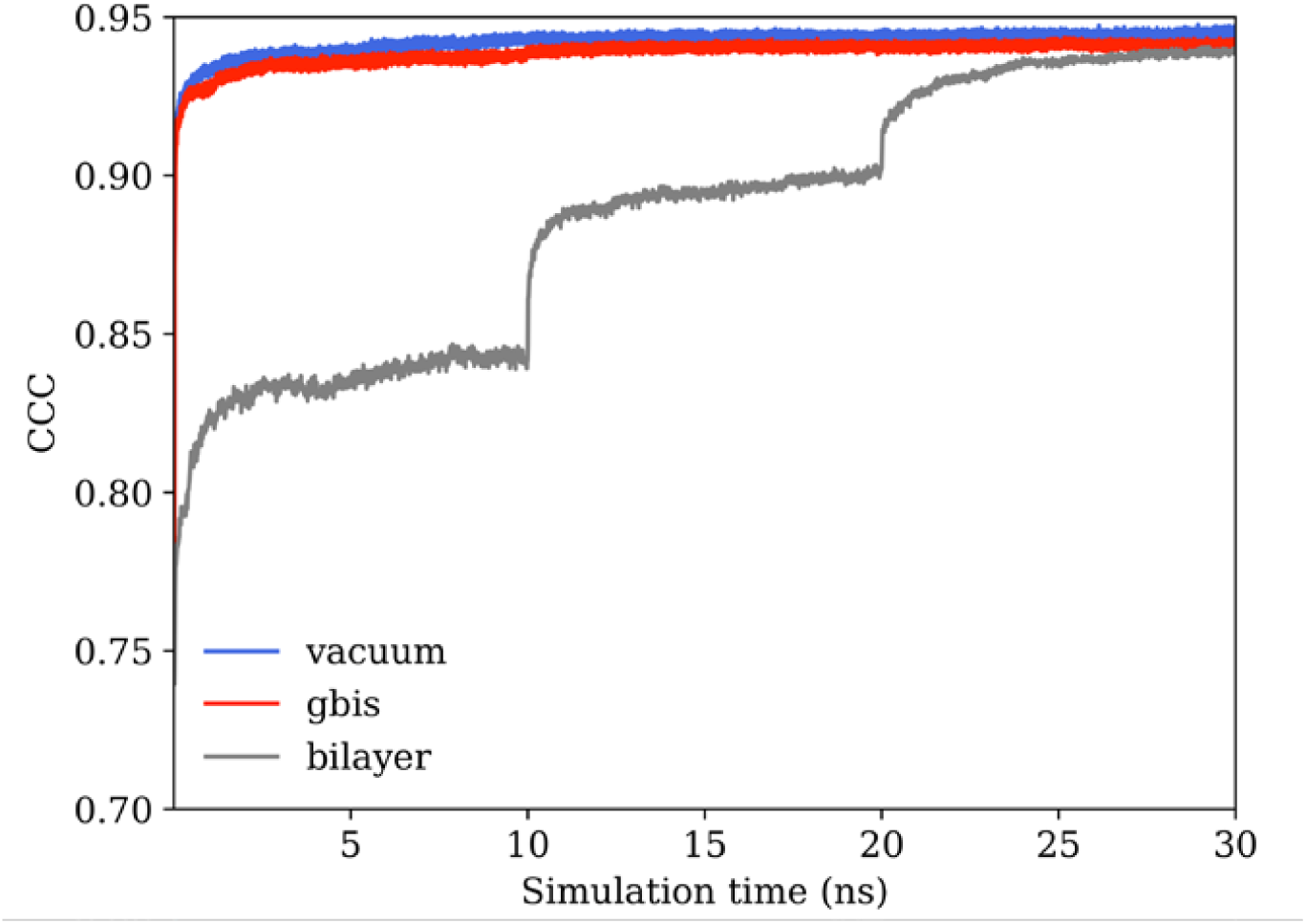
Cross-correlation coefficient (CCC) during the MDFF simulations.

The global measurements of RMSD and CCCs showed little difference in the helical backbone configurations of the hERG fitted structures, particularly in the VSD regions. As a more local measurement, we used MolProbity(37, 38) to check the side chain quality of the models. The MolProbity statistics are reported in Table 1 along with the MolProbity score, which is a single metric to compare overall quality of the models. We also tested the cryo-EM structure. Ramachandan outliers, measuring the backbone phi/psi angles, are most frequent in the vacuum model (3.46%), and least frequent in the gbis model. However, side chain rotamer outliers are most frequent in the gbis model (4.56%) and least common in the cryo-EM model (1.08%). C-beta deviations are most dominant in the gbis model (198), are similar for vacuum and the bilayer model (169 and 167, respectively), and not present in the cryo-EM model. Clash scores show an opposite trend: 11.12 for cryo-EM model, 0 for all the others. RMS deviations of bonds and angles are lowest for the cryo-EM structure, and remain low for all other fitted structures. Finally, the MolProbity score for the overall side chain quality is worst for the cryo-EM model (2.12), and is best for the bilayer fitted MDFF model (1.32). Vacuum and the gbis fitted models perform almost similarly (1.52 vs 1.50). In light of these side-chain quality analyses, it seems that the bilayer fitted MDFF structure quality is superior to other counterparts, including the published cryo-EM structure. This is not surprising as even high-resolution cryo-EM structures often contain poor side chain conformations and interatomic clashes(42).

**Table 1.** MolProbity statistics for cryo-EM and MDFF fitted structures of hERG using different environmental conditions. Analyses are carried out on the entire tetramer. The MolProbity score is shown in *italic*.

| MolProbity statistics |  |  |  |  |
| --- | --- | --- | --- | --- |
|  | Cryo-EM | vacuum | gbis | bilayer |
| Ramachandran outliers (%) | 1.65 | 3.46 | 1.08 | 2.27 |
| Ramachandran favoured (%) | 90.66 | 89.85 | 91.58 | 91.63 |
| Rotamer outliers (%) | 1.08 | 4.14 | 4.56 | 2.62 |
| C-beta deviations | 0 | 169 | 198 | 167 |
| Clash score | 11.12 | 0.00 | 0.00 | 0.00 |
| RMS(bonds) | 0.01 | 0.04 | 0.03 | 0.03 |
| RMS(angles) | 0.99 | 3.52 | 3.53 | 3.57 |
| *MolProbity score* | *2.12* | *1.52* | *1.50* | *1.32* |

### 3.2 Refinement of state-dependent interactions in the VSD establishes additional salt bridges

After assessing the overall backbone and side chain properties in previous sections, we now turn to the refinement of state-specific interactions in the MDFF-fitted hERG models, in particular the presence of functionally important salt bridges in the VSD. A summary of all intra-domain salt bridges in the VSD is presented in Table S1. We analyzed all four chains separately to check the symmetry of the interactions, which may shed light on the activation dynamics of the channel. The cryo-EM structure contains three salt bridges (D411-R541, D460-R528 and D501-R537) and these salt bridges are present in all four chains in the symmetric tetramer. Several other physiologically relevant salt bridges in the depolarized VSD are absent in the cryo-EM structure (3, 6, 8, 11, 15, 18). We now want to explore whether the refinement protocol preserves these three salt bridges from in the cryo-EM structure and further establishes other salt bridges in our fitting procedure. The vacuum MDFF procedure leads to the formation of several additional salt bridges in the VSD of hERG. However, the three pairs present in the cryo-EM are lost in some chains. For example, D411-K538 is lost in the B and D chains and D501-R537 in the A and D chains. The D460-R528 pair is lost chains except B, C, and D. Among newly formed salt bridges, only D466-K538 is present in more than one chain. Hence, the vacuum fitting protocol may not be sufficient to further refine the hERG structure and establish the missing interactions

The gbis and bilayer MDFF procedures lead to the formation of more functionally relevant salt bridges (Table S1 and Figure 3). All salt bridge pairs formed by D411 are present in both cases. However, both procedures failed to produce the D411-K538 in chain D in comparison to the cryo-EM structure. D456-R528 pairs are present in all the chains in the gbis structure and are missing in chains A and D in the bilayer structure. However, R528 forms a salt bridge with D509 in the bilayer structure. D509 is known to be important in hERG function(18). The salt bridge D460-R528 is present in all the chains in the bilayer structure, however are missing in chain D in the gbis structure. The salt bridge D460-R531 is thought to be present in the open state(15). D466 establishes multiple salt bridges with R537, K538, R534 and K407 in the gbis model and to a lesser extent in the bilayer model (only with K538 and R534). The D501-R537 salt bridges present in the cryo-EM structure are retained fully in the gbis model, however this salt bridge is missing in chain D for the bilayer model. In both the gbis and bilayer model, D501-R534 forms in all chains except chain D. The D509-R528 salt bridge, crucially important for activation and stabilization of the open state of hERG(10, 18), is established in chain A and D in the presence of bilayer as stated above. We also see the formation of several other glutamate pairs in both the gbis and the bilayer fitted structures, which do not occur in the vacuum structure or in the cryo-EM structure. In total, the number of salt bridges formed in the gbis or bilayer fitting protocol (44 and 43, respectively) is almost four times higher than the vacuum protocol or the actual cryo-EM (13 and 12, respectively) structure (Table S1).

**Figure 3.**
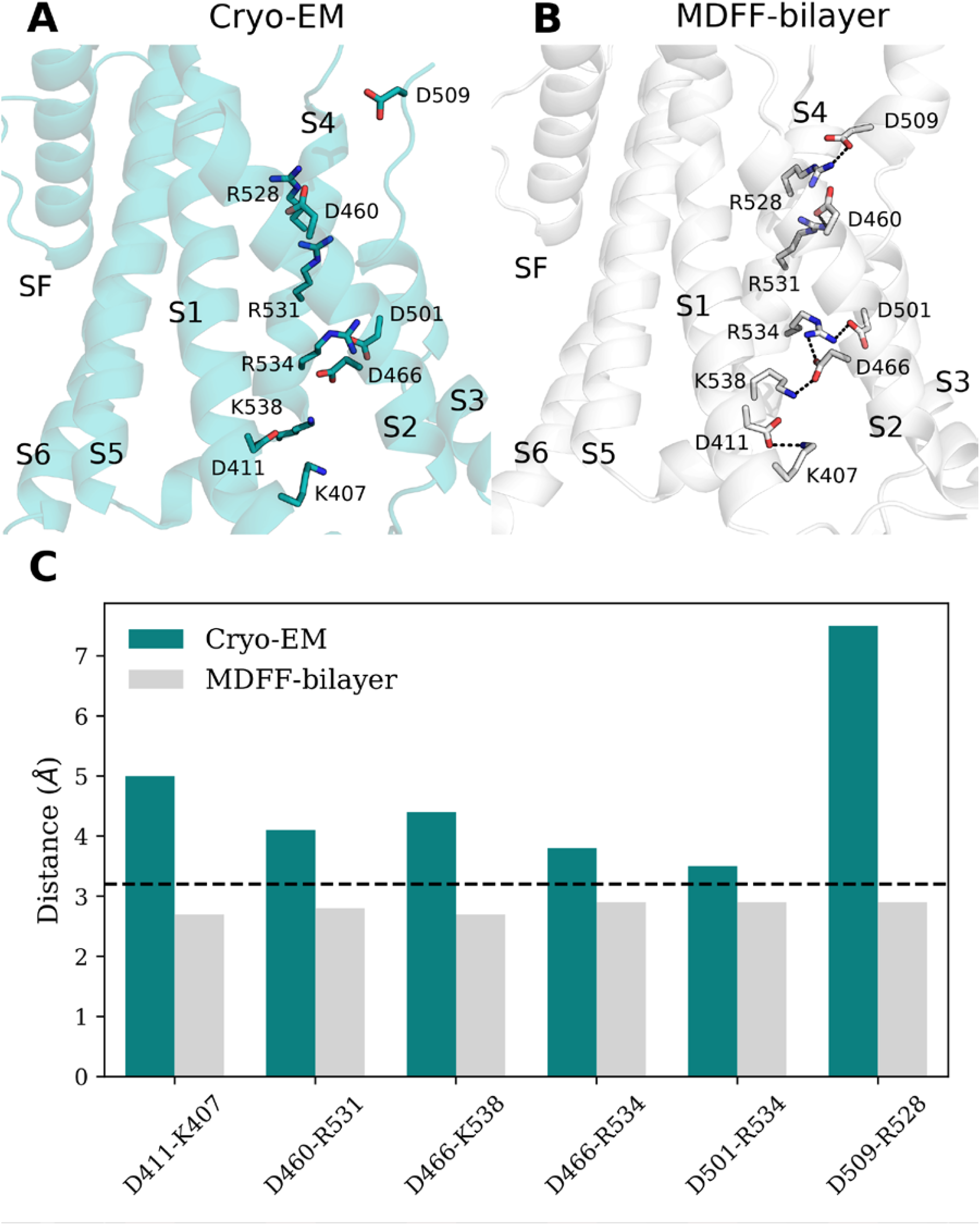
Formation of functionally relevant salt bridges after fitting. (A) Cryo-EM structure for comparison, (B) structure obtained from MDFF-bilayer fitting. Only newly formed salt bridges involving aspartates in chain A are shown for clarity. Dashed lines in (B) represent formation of the salt bridges. (C) Distance between nitrogen of arginine/lysine and oxygen of aspartate in the corresponding pairs.

### 3.3 A novel salt-bridge predicted in the VSD of hERG is important functionally

The MDFF fitting with cryo-EM maps resulted in the formation of a novel salt bridge interaction in hERG VSD formed by D501-R534. To investigate a possible functional role of this salt bridge in activation, we performed electrophysiology experiments (Figure 4) of single and double mutants of the channel. Interestingly, the single R534E mutation elicits a functional hERG-like channel but with altered activation kinetics (Figure 4(B)). These altered functions include a shift of the V_1/2_ of activation to more depolarized potentials (Figure 4(E)) and the development of a slow component of deactivation. The slow tau of deactivation increases from 299 msec in wild type channels to 500 msec in the R534E currents. These features could be interpreted in terms of changes in the barrier associated with the voltage sensor movement during activation process. The shift to longer tau of deactivation in R534E suggests an increase in energy penalty associated with the activation gate opening, but once the gate is open it allows WT-like currents. Interestingly, the R534E mutation has only modest impact on the channel recovery from inactivation. The reversal potential for R534E is -78 mV and the inward rectification and tail currents features associated with recovery from inactivation are comparable to that observed in the wild-type hERG. Importantly, the D501K channel does not conduct any functional current (Figure 4(C)).

**Figure 4.**
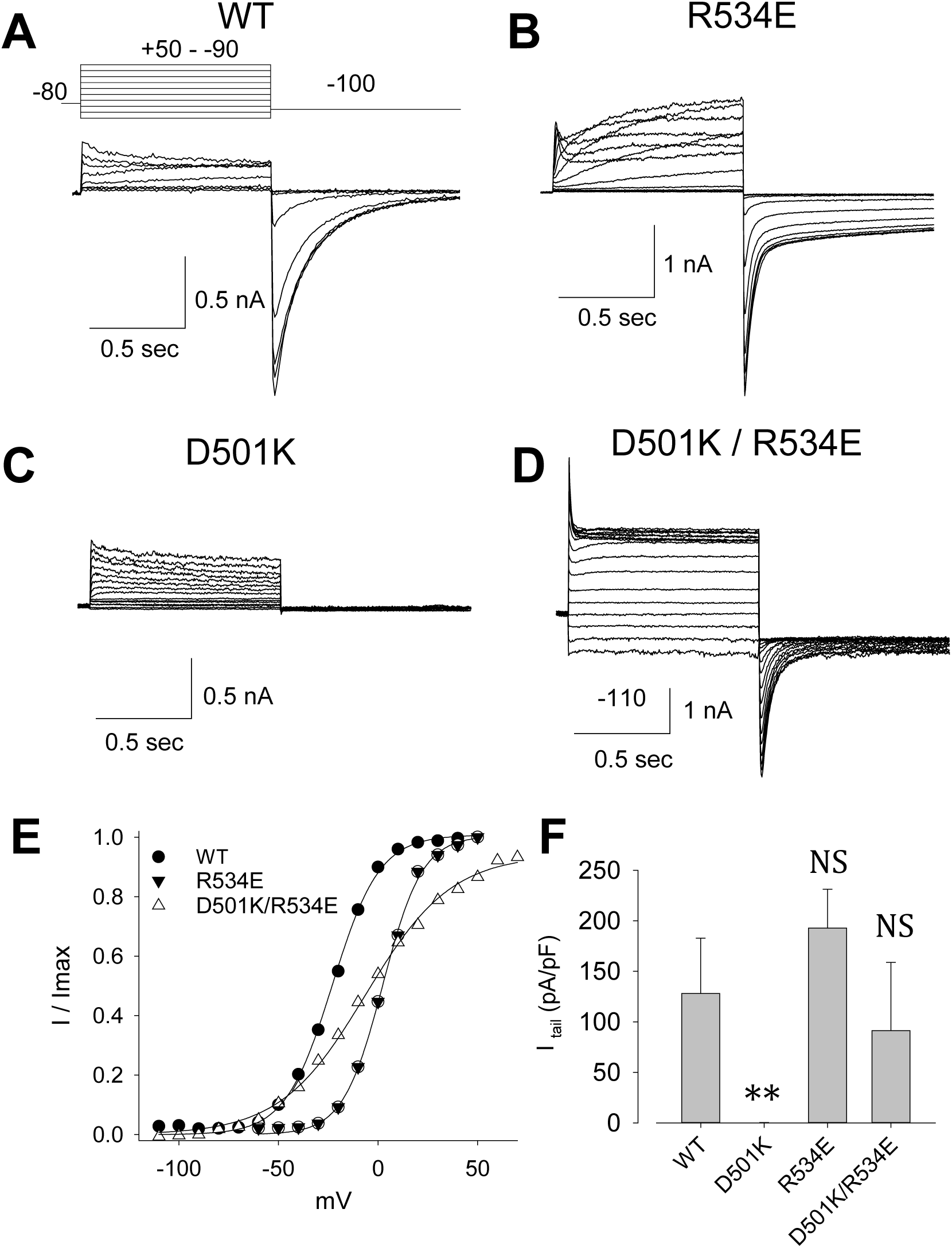
Representative current traces of WT (A), R534E (B), D501K (C) and D501K/R534E (D). D501K did not show visible current. I-V relationships of the tail currents were showed in E. The experimental protocol shows in inset of A. F, The current densities of maximum tail current of each mutation. N= 18, 5, 4, 20 in WT, D501K, R534E, D501K/R534E respectively. NS, no significance; ** P<0.001 comparing to WT in one way ANOVA test.

To further test presence of the salt bridge proposed from MDFF modelling we performed the salt bridge reversal mutation D501K/R534E. The double mutant rescues a functional current expression (Figure 4(D)) but with altered function providing direct evidence for a potential importance of the salt-bridge spatial organization (3, 15, 43). Its alterations include: 1) near instantaneous leak conductance with has a normal reversal potential and 2) tail currents that show a markedly abnormal slope factor for activation. The leak conductance with WT-like reversal potential may reflect presence of the partially opened activation gate in the pore domain in all gating states of the voltage-sensor. At the same time the ion flux occurs through the pore domain and WT-like selectivity filter, allowing WT-like potassium selectivity and C-type inactivation features. During repolarization, a very slow deactivation component is observed with a persistently open leak component, in keeping with an activation gate in the partially open state. We hypothesize that the change in the slope factor for gating and the leak current may reflect abnormalities in the co-operative transitions on the path to channel closing.

## 4. Discussion

The presence of negative charges in S1-S3 helices and positive charges in S1 and S4 helices naturally raises the question of their role in gating processes. Although the direct modelling of the gating process is outside the scope of this study, we can use extensive functional studies published to date to first assess state of the sensor captured in the original cryo-EM structure and then connect to fitting protocols as well as VSD plasticity features emerging from simulation studies. Here we focus our discussion to the interactions present in the bilayer MDFF structure, as this structure seems to be the best in our structure quality checks as outlined in the results sections.

### 4.1 Structure-function relationship is established in the VSD of the refined structure for open-state of hERG

In the cryo-EM structure, only three pairs of salt bridges are present (Table S1). They are: D411-K538, D460-R528, and D501-R537. Hence, there is a clear gap between the structure of this “open-state” resolved by cryo-EM and the other previous functional studies. In the following paragraphs, we discuss the additional salt bridges that were observed in our refined structure and their functional relevance in open, close or transition state stabilizations based on previous works.

In a series of papers, Tseng and co-workers highlighted the importance of these salt bridges in stabilizing hERG functional states and its gating process (5), reviewed by Vandenberg et al(1). Here we focus our discussion on the interactions present in the bilayer MDFF structure (Figure 3). The negatively charged residues form essential interactions as found by Liu et al. (3). In charge neutralization experiments, D460 and D509 form salt bridge characteristic to the open state of hERG1. However, the authors could not determine the positively charged counter-parts for these salt bridges in those experiments(3). We also see bi-furcating salt bridges involving D411 (D411-K538, D411-K407), which are thought to stabilize the closed state of hERG by Liu et al(3). This is surprising as we start from the putative open state of hERG from the cryo-EM structure, yet we see salt bridge formation in our fitted model by a negatively charged residue which is supposed to stabilize the closed state(3-5). In a later charge reversal mutagenesis analysis by Zhang et al., it was postulated that D411 might be involved in stabilizing “multiple gating states” (4), which supports the presence of salt bridges involving D411 in the putatively open (depolarized) structure emerging from MDFF studies.

We also see the formation of the D411-K407 and D411-K538 pairs. We see the simultaneous presence of D411-K407, D411-K538 and D456-R528 pairs and not the D456-K525 pair (Table S1). However, we cannot investigate the functional couplings of these pairs from our simulation alone. We also see that R531, R534 and R528 form salt bridges; they have been implicated to pair with negative charges in the open state by interacting with D411 and D456(4). The importance of R531 has been tested by Piper et al. using double mutant cycle experiments(15). Based on their work and work on the homologous Kv1.2 structure, the authors suggested a functionally important interaction between R531 and D456 to stabilize the open conformation of the channel; cooperative interactions between R531 and D456, D460 and D509 in inactivation process (15). Although we do not see R531-D456 pair, we do find the persistent salt-bridge formation between R531 with D460. Using tryptophan scanning mutagenesis, Subbiah et al. suggested the salt bridge pairing between R528 and R531 in different conformational states; R528 being involved in “at least one closed state and a transition state”, while R531 being involved in “at least one closed and one open state”(11). Our salt bridge analysis is in accord with their observation. D509 has been identified to be very important in stabilizing both the activated and the relaxed state of the VSD and proposed to be directly interacting with a basic residue (R528) or water mediated electrostatic interaction with a basic residue (K525) in S4(18). We also observe the interaction of D509 with R528, however only in the two chains. Several salt bridges involving glutamate are also present in our fitted structure.

Based on these comparisons, we find that the cryo-EM structure is at best a transition state and does not represent a fully functional open state of hERG as many relevant functional interactions are missing. On the other hand, the refined structure represents a functional open state as supported by literature work and our electrophysiology experiments.

### 4.2 VSD in open-state of hERG can be less plastic, tightly packed and is still hydrated by water

One of the important features of the VSD in non-swapped topology channels is its relatively loose packing, which might be important functionally(3, 11, 18). That is, representing, important ensemble of states for protein domain with significant plasticity and linking it to its functional manifestation may be particularly challenging for these channels. The initial structure we started with for fitting was derived from long microsecond MD simulations and the VSD was loosely packed and showed average conformation with RMSD to cryo-EM state of ∼6Å (Figure 1). During the course of fitting, this loosely packed VSD fits into the cryo-EM density map. Hence, the S4 containing VSD is tightly packed in our final structure even in the presence of lipid during fitting (Figure 5).

**Figure 5.**
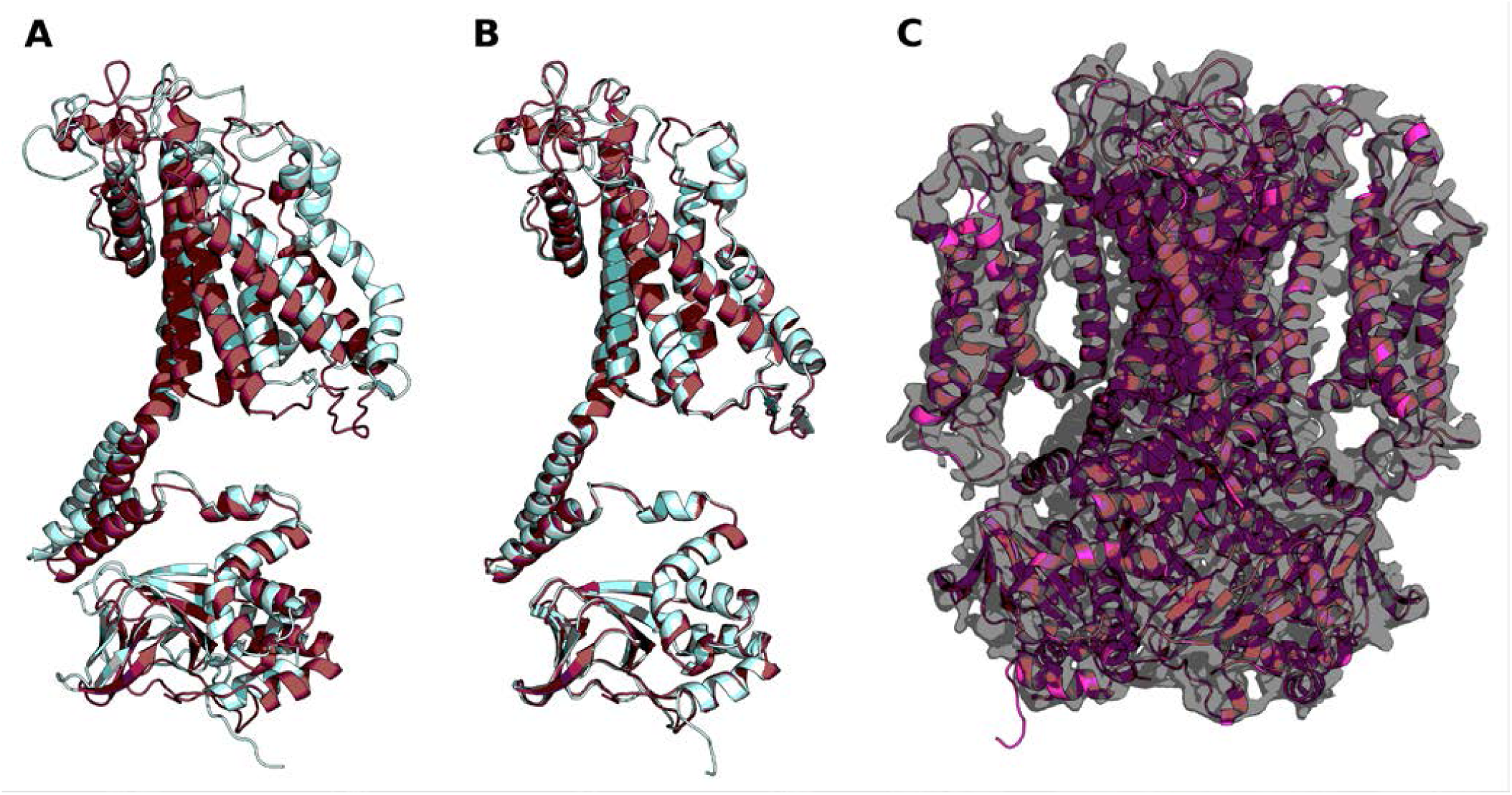
MDFF simulation with POPC bilayer. (A) Structures before fitting, (B) structures after fitting. For clarity, only chain A is presented. Both the initial (pale cyan) and the target (raspberry) structures are presented in cartoon representation. (C) Final structure after MDFF simulation in bilayer and corresponding target cryo-EM map.

We observed a compact configuration of VSD regardless of the environment during the fitting. This is different from the observation on the voltage-sensor flexibility found in a micro-seconds long MD simulation with the reported RMSDs reaching a plateau at around 5 Å(25). A study by Miranda et al. was centered at the permeation across the PD and employed a strong biasing electrical field (750 mV) to enable ion movement which may impact VSD conformation(25). Applications of electrical field are known to induce significant conformational plasticity of VSD in the previously studied Kv channels(44).

The authors observed the rapid relaxation of the VSD in the presence of lipids during MD simulations with an increase in VSD hydration, and subsequent deviations from the cryo-EM structure reported by Wang et al.(16). Interestingly, we see formation of water wire even in this compact VSD in our bilayer fitting (Figure 6). Although, the number of water molecules are significantly less than what has been observed in the unbiased simulation where the VSD adopts an expanded configuration and allows for more water molecules to hydrate the VSD (Unpublished data). As the VSD is constrained by the density map in our fitting procedure, it is natural that the VSD will have a tightly packed configuration (resembling the resolved cryo-EM state and deviating less from it) and has smaller hydration numbers. Hydration of the VSD observed in the fitting process highlights the importance of the VSD plasticity for hERG function, which, albeit established experimentally, has been largely overlooked so far in the channel modeling/structural data interpretation.

**Figure 6.**
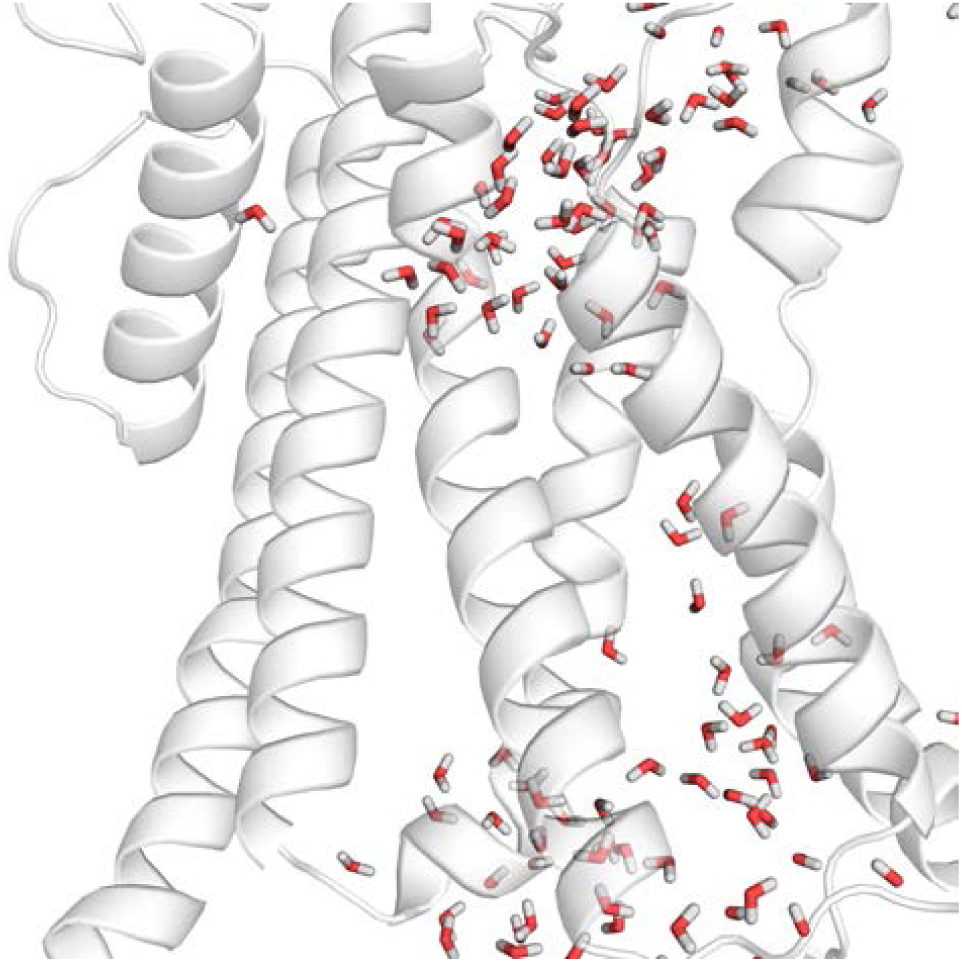
Formations of water wire in the VSD during bilayer fitting highlighting VSD plasticity.

## 5. Conclusions

By using MDFF, we manage to further refine the hERG open state cryo-EM structure. Our bilayer fitted MDFF structure seems to be the most relevant open state representation of the hERG considering the global structural quality, side chain model quality and the functionally relevant salt bridges. We also identified the presence of an additional novel salt bridge in the VSD, which has been further confirmed with electrophysiology experiments. Our integrated approach leads to a fully functional open state hERG structure, and can be used for other ion-channels and transporters.

## Author Contributions

H.M.K. designed and performed the MDFF/MD simulations, analysis of Cryo-EM maps and resulting data analysis. J.Q. designed and performed all molecular-biology and electrophysiology experiments to test novel salt-bridge predicted by the MDFF refinements. S.Y.N., D.P.T and H.J.D. conceptualized the study, designed experiments, and interpreted results. H.M.K. wrote the first draft of manuscript. All authors wrote, edited, and contributed intellectually in writing the manuscript and interpreting the data.

## Acknowledgements

Work in S.Y.N.’s group was partially supported by NIH Grant (R01HL128537-03); this work was supported by the Canadian Institutes for Health Research (Project Program Grant FRN-CIHR 156236, S.Y.N., D.P.T. and H.J.D). D.P.T. acknowledges support from the Canada Research Chairs program. All calculations were performed on the CFI/NSERC-RTI-supported GlaDos cluster at the University of Calgary and on the West-Grid/Compute Canada clusters under Research Allocation Awards to S.Y.N. and D.P.T. H.M.K. acknowledges funding from the University of Calgary through the “Eyes High Postdoctoral Fellowship” program. H.M.K. would like to thank Meruyert Kudaibergenova and Williams Miranda for stimulating discussions about hERG.

